# Molecular Characterization and Phylogenetic analysis of *Wuchereria bancrofti* in human blood samples from Malindi and Tana River Delta, endemic regions in Kenya

**DOI:** 10.1101/648220

**Authors:** Kinyatta Nancy, Wambua Lillian, Mutahi Wilkinson, Mugasa Claire, Kamau Luna, Solomon K. Langat, Wachira Dorcas, Ichugu Christine, Waigi Emily, Githae Rosemary, Lusweti Japheth, Kagai Jim

## Abstract

**Introduction:** Lymphatic filariasis is a debilitating disease caused by filarial worms; *Wuchereria bancrofti, Brugia Malayi* and *B. Timori*. It is earmarked for elimination by the year 2020 through the Global Program for the Elimination of Lymphatic Filariasis (GPELF). In Kenya, mass treatment has been ongoing since the year 2002 though it has not been consistent as recommended by World health organization (WHO). Taking this into account, the emergence of *W. bancrofti* resistance strains against the current choice of drugs cannot be ruled out. Information on genetic structure and variations is important in assessment of Program’s success. Data on genetic characterization of *W. bancrofti* in Kenya is lacking. This study, therefore reports the first genetic diversity of W. *bancrofti* in two Kenyan endemic regions.

**Methodology:** Genomic DNA was extracted from 100 human blood samples obtained from Mpirani district in Malindi and Kipini district in Tana River Delta. They were then amplified by PCR and detected through gel electrophoresis. Seventeen PCR products positive for *Wuchereria* PCR *bancrofti* were purified and then DNA quantified for Sanger sequencing. Chromas version 2.6.5 and BioEdit softwares were used for sequence alignment and editing. Fourteen sequences were selected for analysis by MEGA7 and six more related sequences retrieved from the Gene Bank for further analysis with the study sequences. Intrapopulation, interpopulation diversity and pair wise distance were determined and the phylogenetic trees constructed. Tajima’s D-test of neutrality was also determined and Statistical evolutionary rate was done using Chi-square (X^2^) test.

**Results and Discussion:** The mean diversity of Malindi and Tana River Delta isolates was 1.42 and the overall mean distance was 0.99. Tajima’s (D) test for test of Neutrality was 4.149 and nucleotide diversity(π) was 0.603. These results revealed high genetic variations of *W. bancrofti* in Kenyan endemic regions. This variation could be attributed to prolonged use of the mass drug administration (MDA) and the long period of parasite circulation in these populations.

**Author Summary:** Elephantiasis is a disabling disease that causes severe swellings to the affected limbs. It is caused by parasites of *Wuchereria bancrofti, Brugia Timori* and *B. malayi* which are transmitted by mosquito vectors. The disease is under the control by the Global Programme to eliminate filariasis and due to the effect of continued treatment through mass drug administration there have been changes in the genetic makeup of the parasite. This may result to resistant strains which may have negative impact on the treatment interventions. We therefore aimed at characterizing the genetic sequences of the *Wuchereria bancrofti* parasite found in Kenya. Through analyzing parasites obtained in different years after treatment, we were able to track any genetic variations since the start of mass drug administration in Kenya. These variations would be due to the effect of drug pressure, human population movements or mosquito vector movement. This kind of study is important for drug developments and for evaluating the progress of the control programmes.

## Introduction

Lymphatic filariasis (LF) commonly referred to as elephantiasis is a disfiguring and disabling disease caused by filarial nematode worms. The disease is widespread and a major public health problem in many developing countries with warm and humid climate and is one of the most neglected Tropical diseases causing severe suffering and socio-economic burden in endemic areas. The three lymph dwelling parasites include; *Wuchereria bancrofti, Brugia Malayi* and *B. Timori* which are transmitted by mosquitoes of various species [1], [2]. More than 90% of Lymphatic filariasis is caused by *Wuchereria bancrofti* (Wb) parasites [3]. Current estimates suggest that more than one billion people living in endemic areas are at risk of infections, and more than one third of the infections are in Sub-Saharan Africa [4].

World Health Organization (WHO), in 2000 launched the Global Programme to Eliminate Lymphatic Filariasis (GPELF) with the goal of eliminating filariasis by the year 2020 [5]. The GPELF has two principal pillars: (i) to interrupt LF transmission, and (ii) to manage morbidity and prevent disability. Under this Programme, the principal measure recommended for lymphatic filariasis control is annual community-wide mass drug administration (MDA) of a single dose of 400mg albendazole combined with 6 mg/kg of diethylcarbamizine citrate (DEC) or 400mg/kg Ivermectin to identified communities in endemic areas [6] for 4-6 years with 80% compliance coverage [7]. Progress towards this goal is promising, with 3.9 billion doses of medicine distributed to people in 65 countries by 2011, out of 73 countries considered to be endemic in 2014, 20 had progressed to the post-MDA surveillance phase by 2016 and 52 required further rounds of MDA [8].

In Kenya, LF is prevalent in the coastal region where ecological factors are suitable for transmission [2], [9]. Kenya launched MDA in Kilifi County in 2002. The MDA treatment in Kenya is based on combined single-dose for annual mass treatment in 2003, the Programme scaled up to Malindi and Kwale districts. Two more rounds of MDA were conducted in these districts in 2005, 2008, 2011 and then 2015 [10], [11], [12]. The treatment has however not been consistent with WHO recommendations of annual MDA for 4-6 years which may result in resurgence of transmission and development of genetic variations of the circulating parasites. Additionally, long term chemotherapeutic programs, such as MDA, have been shown to lead to changes in the parasite population structure thus altering management programs and causing potential resurgence of resistance strains after its cessation [13], [14], [15]. The genetic diversity of *W. bancrofti* influences response to drug treatment and parasite fecundity [16] thus having important connotations to the on-going elimination program. Mass drug administration is likely to eliminate susceptible worms, as a result of which the susceptible genes are not passed on to the future generations but only the drug-resistant genes [16]. Monitoring genetic diversity of these parasites thus enables us to understand the response of the parasite population to treatment and allows the identification of resistant strains [17].

Despite the fact that *Wuchereria bancrofti* is a leading cause of disability, very little is available on its genetic structure and diversity. Understanding the genetic variation will be essential for monitoring the success of programs aimed at control and elimination of lymphatic filariasis in regions where *W. bancrofti* is endemic. Until recently, the availability of genetic markers for differentiating between strains of *W. bancrofti* has been limited, so it has not been possible to evaluate changes in parasite populations in the context of LF elimination programs. Through sequencing of the *W. bancrofti* mitochondrial genome (mtGenome) numerous genetic polymorphisms have been identified that can be used to evaluate population structure and to characterize infections. This allows us to move beyond mere detection and track individual *W. bancrofti* strain prevalence through time to time. The length of *W. bancrofti* mitochondria (mt) is approximately 13, 637 nucleotides, contains two ribosomal RNAs (rrns), 22 transfer RNAs (trns), 12 protein –coding genes, and is characterized by a 74.6% AT content [18]. The *W. bancrofti* mt gene order is identical to that reported for *Onchocerca volvulus, Dirofilaria immitis, Setaria digitata* and *Brugia malayi. Wuchereria bancrofti* size length of complete genome sequence is calculated by adding lengths of all scaffolds together 81.51Mbp, 29.70% GC content of scaffolds, 19 327 gene number of predicted protein-coding genes in genome, 112 t RNAs number of predicted tRNA genes in genome and 8 rRNAs number of predicted rRNA genes in the genome [19].

In Kenya, *Wuchereria bancrofti* genetic data is lacking and the current study aimed at molecular sequencing and characterizing of partial *W. bancrofti* 18s rRNA gene from specimens collected from Malindi and Tana River Delta districts where the disease is prevalent. We thus analyzed the Sps1 repeat sequences from two districts with the goal of improving our understanding of the genetic makeup of *W. bancrofti* and determine if there are any genetic variations of *W. bancrofti* parasite within Kenyan endemic regions.

Genetic information of parasites is essential for identifying targets for drug development and the potential of drug resistance. Through genotyping *W. bancrofti* populations we can get information that is likely to contribute to elimination success. The World health organization recommends albendazole, ivermectin and diethyl-Carbamazine for the control of lymphatic filariasis in endemic areas through mass drug administration to at-risk population for 4-6 years. However, the parasites are treated as a single entity chemotherapeutically as well as epidemiologically in the control programs undertaken on a global scale. This generates a factor of ambiguity in the success of such large scale control programs. Hence, the possible development of higher tolerance to the drug of choice for Bancroftian filariasis by some genetic variants may contribute to non-realization of goals of control/elimination programs. It has been evidenced that even after more than 6 years of treatment in some countries, the transmission of the parasite is still on. Continued use of drugs has resulted to resistance strains of *W. bancrofti* especially with albendazole though only few studies have reported drug resistance against *W. bancrofti*.

## Materials and Methods

### Study site

The blood samples for this study were collected from Malindi constituency in Kilifi County and Tana Delta constituency in Tana River County in Coastal Kenya. The study areas were selected because of the high prevalence of LF previously observed which was at 26.3% in Tana River and 22.0% in Malindi [20]. Malindi was among endemic areas where MDA begun in Kenya and Tana River county is among the counties were MDA started later after several rounds in the other endemic areas. Tana River Delta was curved off from Tana River district in October 2007. Tana River Delta has three divisions; Garsen, Kipini and Tarasaa. It has an area of 16,013.4km^2.^ The population in both Tana River and Tana Delta is estimated to be 250,000 with about 134,000 being in Tana River district according to 2009 census [21]. The rainfall ranges between 220-900mm per year and an average temperature of 30°C. The altitude ranges between 0-200m. The major ethnic groups are the Pokomo, many of who are farmers, and the Orma and Wardey, who are predominantly nomadic. Malindi is a town on Malindi Bay at the mouth of the Galana River, lying on the Indian Ocean coast of Kenya. It is 120 kilometres northeast of Mombasa with Coordinates 3°13‵25’40°7’48”E. It covers a geographical area of 7, 605 square kilometers with a total population of 207,253 as of the 2009 census [22] and the main industry is tourist attraction. Malindi forms a municipal council with thirteen wards. It’s a cosmopolitan town with mixed ethnic groups predominant inhabited by Swahili-Arab descendants and Mijikenda including Giriama and Chonyi Communities. The weather is generally warm throughout the year with temperature range above 25°C and two rainy seasons (above 800-1000m). The map of the study area is shown in (figure 1).

**Figure 1;.**
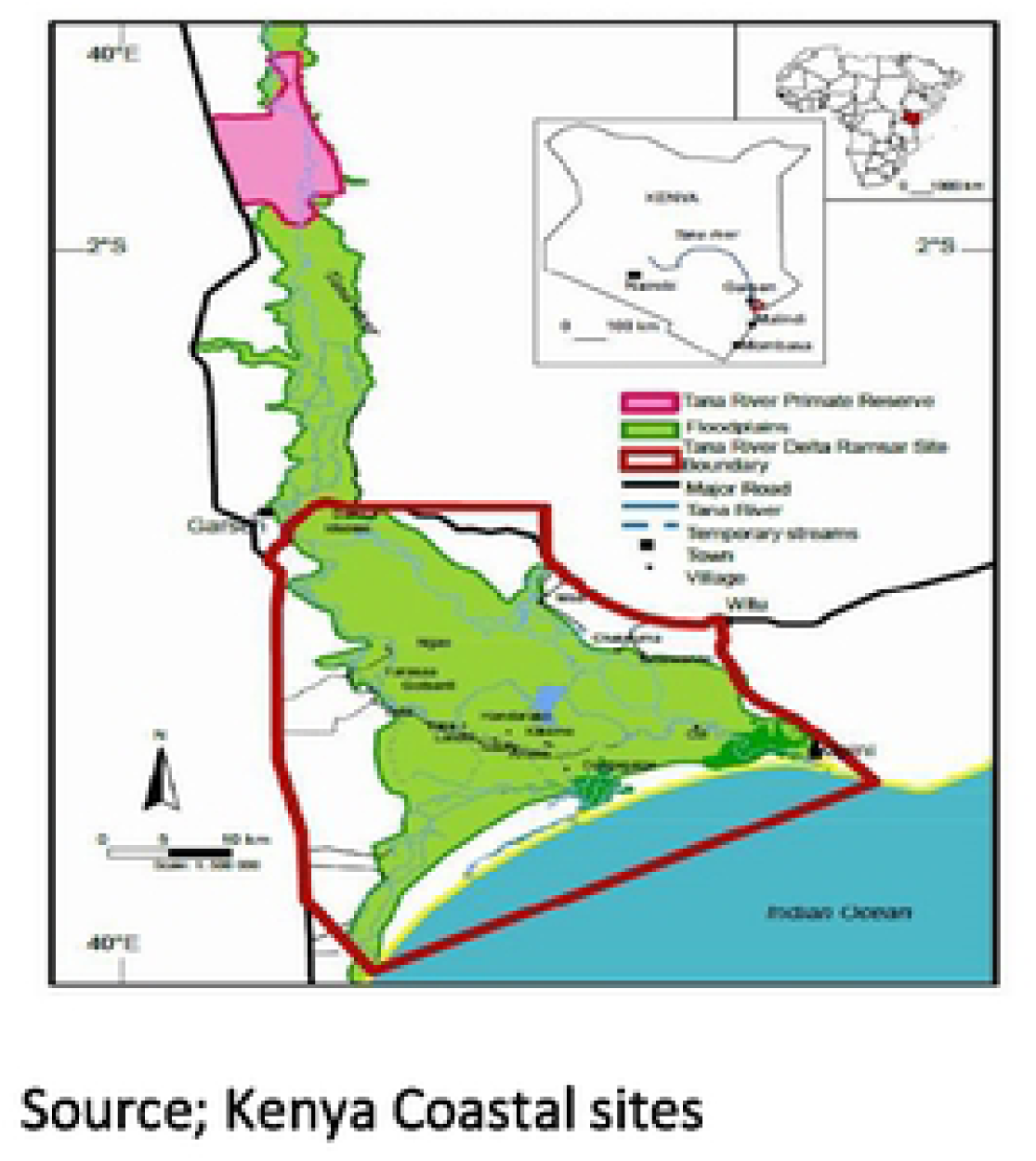
Map of the study area.

### Ethical considerations

Ethical clearance to carry out this study was sought from Kenya Medical Research Institute-Scientific and Ethical research unit (SERU), SSC protocol number 2802.

### Study design

This was a longitudinal retrospective study. For this study, samples were selected from archived samples from previous epidemiological surveillance studies by Kagai and colleagues [20], including Molecular epidemiology of *Wuchereria bancrofti* infections in Tana Delta 2008, The inference of Immunochromatographic test, microscopy and polymerase chain reaction in diagnosis of lymphatic filariasis in Tana Delta, Kenya, 2011 and Molecular technique utilizing sputum for detecting W. b infections in Malindi (2002, 2008, 2011).

Human blood samples before the start of MDA and for subsequent years of MDA were selected from archived samples from previous studies. The samples were collected from Malindi in 2002, 2004, 2011 and from Tana Delta in 2008, 2011. This was to allow for better comparison for detection of the genetic divergence.

### Laboratory Analyses

#### Deoxyribonucleic acid (DNA) extraction from human blood samples

DNA from human blood samples was extracted using the alcohol precipitation method as described by Weil and colleagues [23] with minor modifications. Briefly 200µl of blood was added in to labeled 1.5ml eppendorf tubes and 400µl of sodium hydroxide with 1% triton added. The mixture was heated at 65°C for 30 minutes in a Thermomixer (Thermo Fisher) with continues shaking and quickly cooled on ice. Using a PH meter, the pH was adjusted to 8 for optimal performance of the Taq Polymerase during amplification. The DNA was centrifuged at 1400rpm, 4°C for 5 minutes and the supernatant transferred to a new eppendorf tube. Eight hundred (800) µl of absolute ethanol (98%) was added, vortexed and incubated overnight at - 80°C. The DNA was then washed thrice with 70% alcohol, dried and 50µl Tris-EDTA (TE) buffer added and stored at −20°C until amplification was performed.

#### Amplification of *W. bancrofti* 18S rRNA gene by Polymerase Chain Reaction

Polymerase chain reaction (PCR) was performed using NVI (5’ CAACCAGAATACCATTCATCC 3’) as the forward primer and NV2 (5’CGTGATGGCATAAAGTAGCG 3’) as the reverse primer as previously described by Zhong [24] and WHO, [25]. The target sequence for these primers is the species specific (Ssp 1) repeat sequence of the 18s rRNA gene present at ~ 500 copies per haploid genome. Amplification with these primers yields 188bp fragment. A 25µl PCR reaction was prepared containing 10x Bioline buffer (with Mgcl_2_ and dNTPs), 5pmol/µl of NV1 and NV2 primers each, 5µl genomic DNA template and water to top up the reaction volume. The PCR reaction was run in a 96-well Gene Amp® PCR system 9700. The reaction conditions consisted of a single step of 95°C for 5 minutes, 40 cycling step of 94°C for 30 seconds, 54 °C for 45 seconds, 72 °C for 30 seconds and a final extension step of 72°C for 10 minutes. The PCR products were size fractioned on 2.0% agarose gel stained with Ethidium bromide. Agarose gel electrophoresis was carried out at 80V for 60 minutes and bands visualized under UV light using a gel documentation system (EZ Imager, Bio Red, CA). Positive control used was generously provided to Jim Kagai by Hamburger, Hebrew university, Israel and negative control used was PCR water and blood sample from filarial non-endemic area to ensure specificity and validity of the results. The expected *W. bancrofti* size band was 188 base pairs and this was measured against a 100 base pairs molecular weight marker.

#### PCR product purification, quantification and sequencing

Detected amplicons were purified using QIAquick protocol spin columns in a micro centrifuge as per the manufacturer’s manual. The DNA quality and quantity concentration measurement was done using Nanodrop 2000 (Thermo Fischer Scientific), one (1) µl of the concentrated products was used and the concentrations ranged from 90.2ng/µl to 30.0 ng/µl. The DNA concentration was adjusted to 50ng/µl by adding TE buffer. 2µl of the concentrated DNA were analyzed in gel electrophoresis. Five microliters of each of the selected amplified products were sent for Sanger sequencing (Macrogen-Europe).

#### Molecular Characterization of *W. bancrofti* based on the Ssp1 DNA repeat sequence

The sequences imported into and assembled Chromas software Version 2.6.5, then trimmed using Bio edit Programme. They were aligned using ClustalW with MEGA version7 software. Short sequences were excluded from further analysis. Basic Local Alignment Search Tool (BLASTn) tool was used to search for other related sequences from Genbank nucleotide database (National Centre for Biotechnology Information, NCBI) for homologous gene sequences with that obtained from this study. Genetic characteristics analysis was done by constructing phylogenetic tree and evolutionary distance matrix calculated. The nucleotide sequences for this study have been submitted to Genbank and allocated Accession numbers MK471341 - MK471350.

## Results

### DNA amplification

The PCR products after amplification and detection by gel electrophoresis are shown in figure (2). The *wuchereria bancrofti* product size was 188bp againest 100bp Molecular Ladder for the Ssp I repeat sequence.

**Figure 2;.**
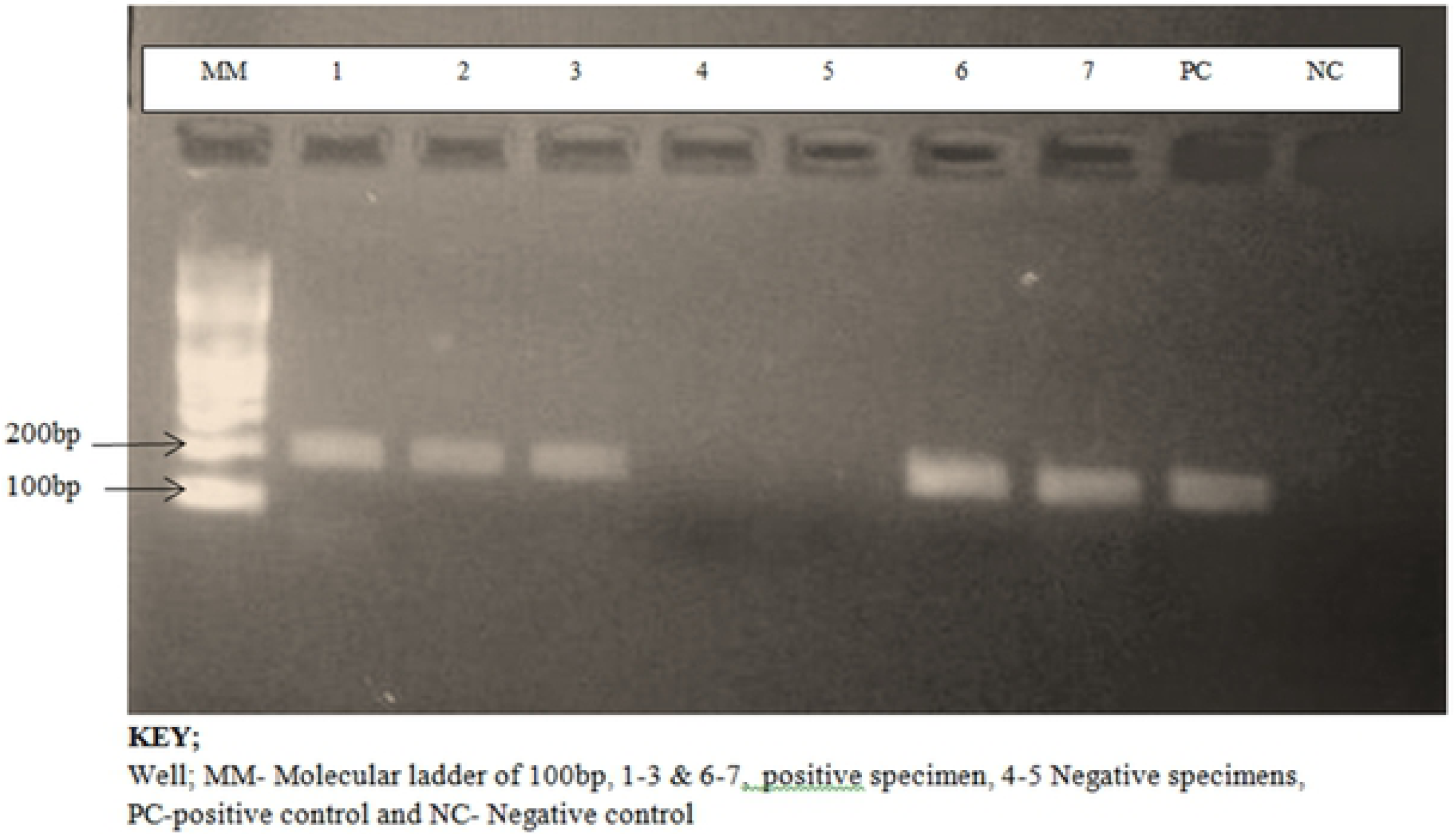
Detection of *Wuchereria bancrofti* DNA on Gel electrophoresis.

### Sequence Analysis

On assembling and trimming of the 17 samples sequenced, only 14 sequences were selected for further analysis. Blast analysis on NCBI showed that most of the sequences were related to sequences belonging to *W. bancrofti* strains on accession numbers; LM012589.1, LM000927.1, AY297458.1, L20344.1, AP017705.1 (table 1) with identity range of 79-98%.

**Table 1;.** Sequenced samples and Gene Bank Blast results.

| Sample ID | Gene description | E-value | % Identity | Accession No. |
| --- | --- | --- | --- | --- |
| ML1 | W. b genomic assembly W. b Jakarta Scaffold WBA contig 0011217 | 8e-44 | 98 | LM012589.1 |
| ML3 | W. b genomic assembly W. b Jakarta Scaffold WBA contig 0011217 | 8e-44 | 79 | LM012589.1 |
|  | W. b genomic assembly W. b Jakarta Scaffold WBA contig 0000579 | 3e-71 | 92 | LM012589.1 |
|  | W. b nuclear Scaffold/matrix, attached region | 3e-71 | 92 | AY297458.1 |
| ML4 | W. b genomic assembly W. b Jakarta Scaffold WBA contig 0011217 | 5e-34 | 95 | LM012589.1 |
|  | W.b nuclear Scaffold/matrix, attached region | 1e-143 | 82 | AY297458.1 |
| ML5 | W. b genomic assembly W. b Jakarta Scaffold WBA contig 0011217 | 1e-05 | 85 | LM012589.1 |
|  | W.b nuclear Scaffold/matrix, attached region | 4e-36 | 83 | AY297458.1 |
| ML11 | W. b Ssp I repeat DNA sequence | 1e-58 | 98 | L20344.1 |
|  | W.b nuclear Scaffold/matrix, attached region | 4e-58 | 98 | AY297458.1 |
|  | W. b mitochondrial DNA complete sequences | 2e-17 | 87 | AP017705.1 |
| TR1 | W. b nuclear Scaffold/matrix, attached region | 1e-48 | 93 | AY297458.1 |
|  | W. b Ssp I repeat DNA sequence | 5e-47 | 94 | L20344.1 |
| TR4 | W. b nuclear Scaffold/matrix, attached region | 1e-45 | 93 | AY297458.1 |
|  | W. b Ssp I repeat DNA sequence | 6e-46 | 94 | L20344.1 |
| TR6 | W. b nuclear Scaffold/matrix, attached region | 2e-52 | 92 | AY297458.1 |
|  | W. b Ssp I repeat DNA sequence | 4e-48 | 91 | L20344.1 |

**Label;** ML- Malindi isolates, TR -Tana River Isolate

### Estimates of Evolutionary Divergence between Sequences of *Wuchereria bancrofti* isolates of Malindi and Tana River

The mean evolutional diversity within the two populations was 1.42, Malindi had the greatest genetic divergence of 1.81 among the isolates of the three sampling times and Tana River Delta had the least genetic diversity of 0.40 for the isolates of the two sampling times. The mean evolutionary diversity of the entire population was 1.98, with Malindi having 2.26 which was higher compared to Tana River Delta which had 0.45. The inter population diversity was 0.56 for both population, Malindi had a lower inter population diversity of 0.46 and Tana River had the least 0.04. The coefficient difference of both populations was 0.28, Malindi had a lower coefficient of 0.2 and the least was 0.1 from Tana River (figure 3).

**Figure 3.**
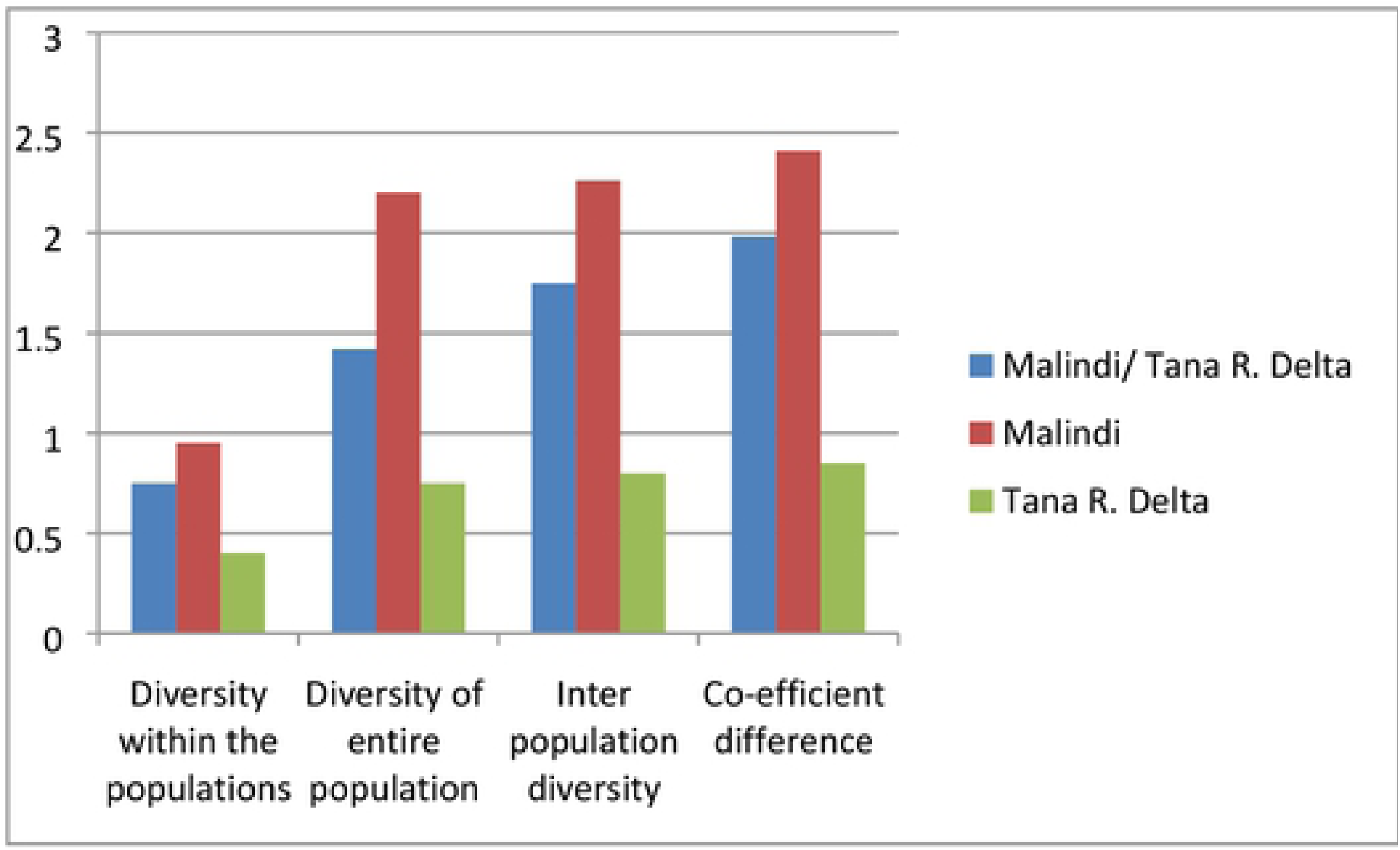
Evolutionary Divergence of Malindi and Tana River Delta Wuchereria Bancrofti Isolates, Kenya.

### Paired wise distance and population mean distance estimation

The Paired wise distance of the population is shown in Table 2. The Overall mean composition distance between the samples was 0.99 and that of within group mean distance was 1.11 and 0.39 for Malindi and Tana River Delta respectively.

**Table 2;.**
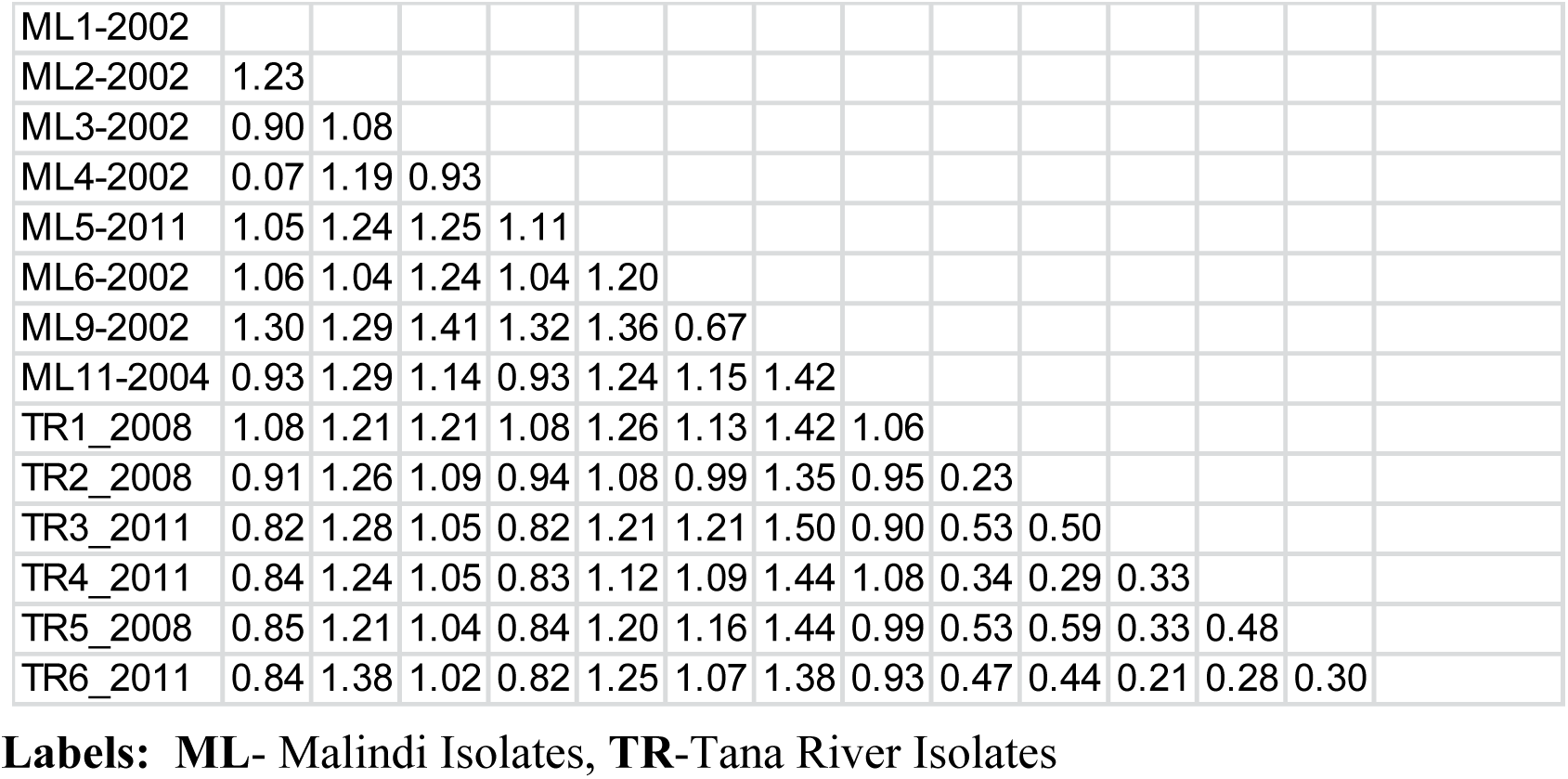
Estimating Evolutionary distance using Paired wise distance.

Analyses were conducted using the Maximum Composite Likelihood model [26]. The analysis involved 14 nucleotide sequences. Codon positions included were 1st+2nd+3rd+Noncoding. All positions containing gaps and missing data were eliminated. There were a total of 262 positions in the final dataset.

### Molecular and Phylogenetic analysis of Malindi, Tana River Delta Isolates and related Genbank sequences

Evolutionary analyses were conducted in MEGA7 [27]. Eight (8) sequences from Malindi and 6 sequences from Tana River Delta isolates were used for phylogenetic tree reconstruction (figure 4). Six (6) more sequences of filarial origin including; *Wuchereria bancrofti* isolate Wb1 07 18S rRNA from Brazil Accession number EU272178.1, *Wuchereria bancrofti* ribosomal protein S13 from Atlanta Accession Numbers; M86642.1, *Mansonella perstans* 18S rRNA from Spain Accession number DQ995498.1, *Loa loa* 18S rRNA from Spain Accession number DQ995497.1, *Wuchereria bancrofti* ribosomal protein S13 from Atlanta Accession number M86643.1 and *Brugia pahangi* mRNA from Atlanta Accession number x16591.1 were retrieved from genbank, aligned and trimmed for evolutionary comparison with the sequences of this study.

**Figure 4.**
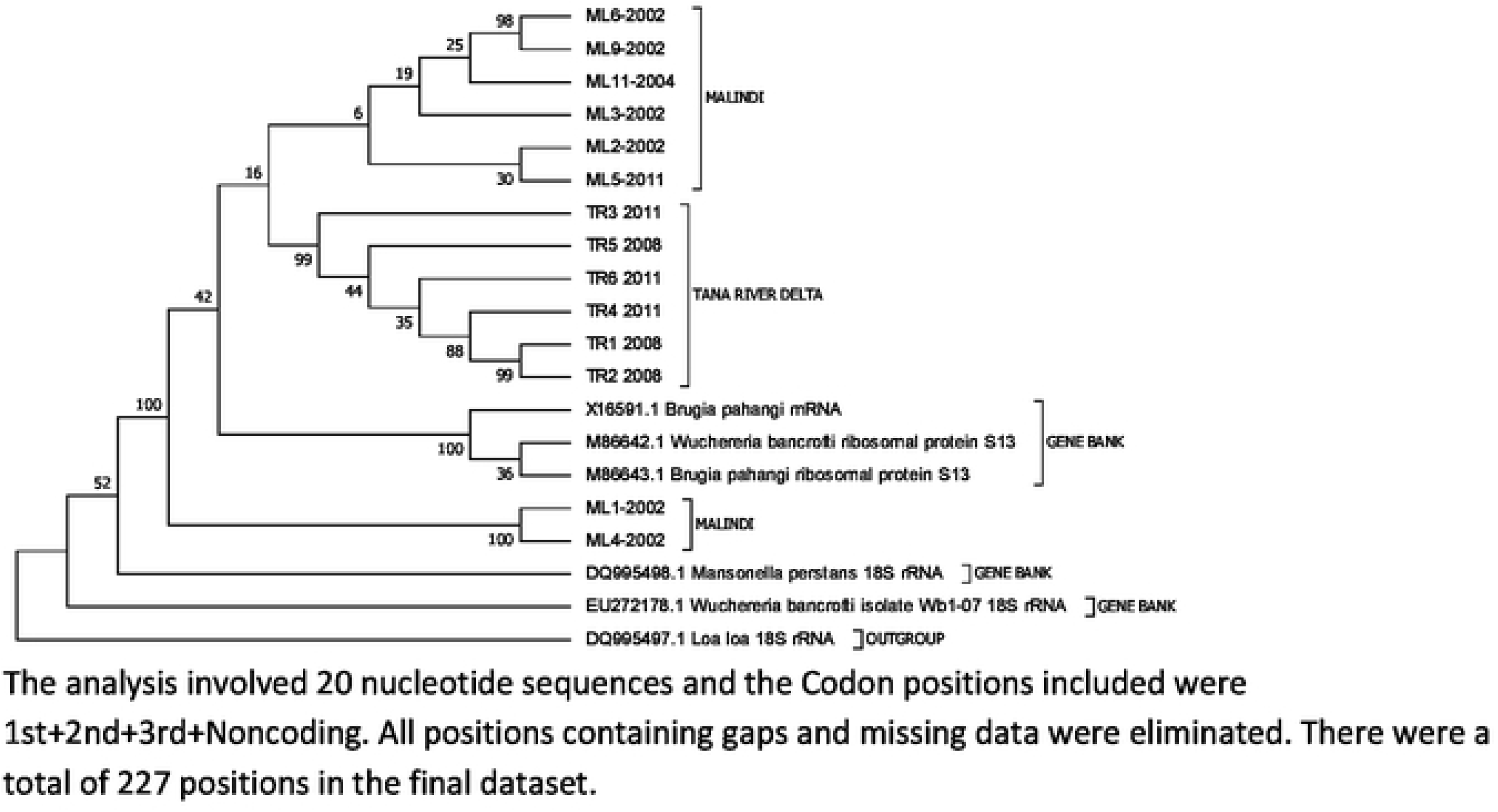
Molecular Phylogenetic analysis of Malindi and Tana River Delta Isolates.

The evolutionary history of the isolates was inferred using the Maximum Parsimony method. The length of the parsimonious tree is 1226. The consistency index is 0.488962, the retention index is 0.572211, and the composite index is 0.279789 for all sites and parsimony-informative sites. The associated taxa clustered together at 1000 replicates bootstrap shown next to the branches as described by Felsenstein [28]. The Maximum Parsimony tree was obtained using the Sub tree-Pruning-Regrafting (SPR) algorithm as per Nei’s manual [29]. The analysis involved 20 nucleotide sequences and the Codon positions included were 1st+2nd+3rd+Noncoding. All positions containing gaps and missing data were eliminated. There were a total of 227 positions in the final dataset. Accession numbers allocated to the nucleotide sequences form this study submitted to gen bank are shown in (Table 3).

**Table 3;.**
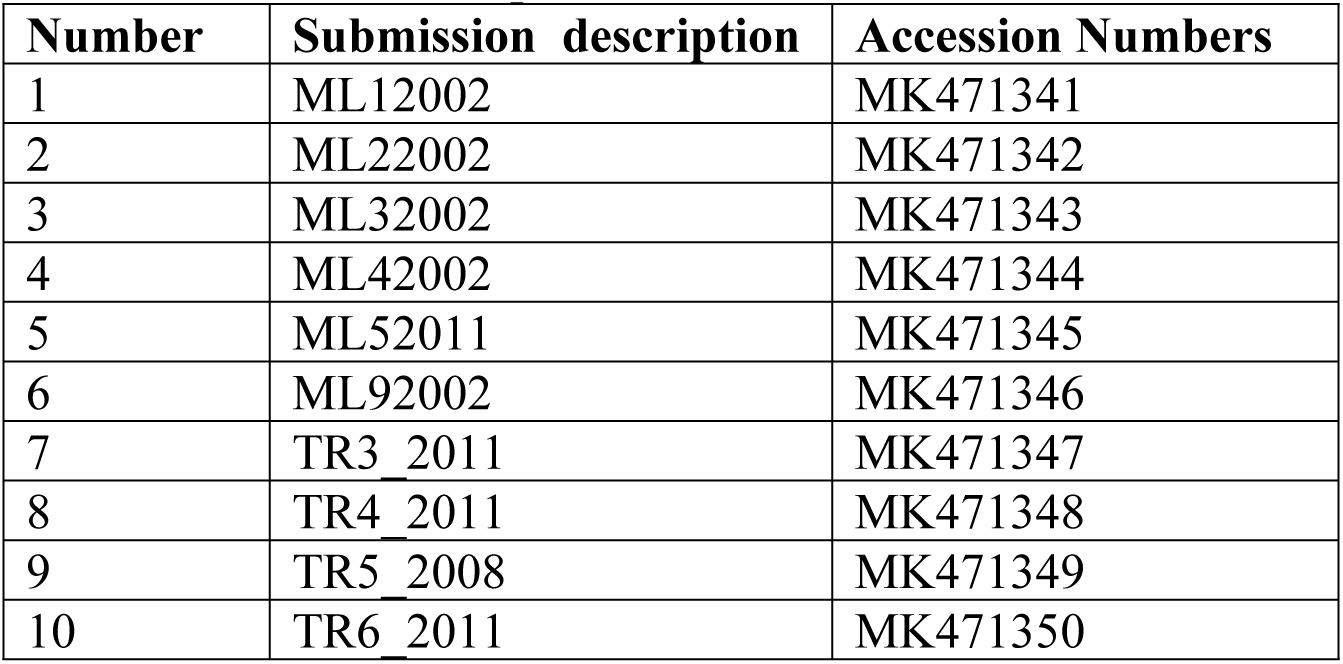
Nucleotide sequences Accession Numbers.

### Neutrality and selection test

In this study a test for selection reinforced neutrality within the Ssp 1 repeat sequence data was done by Tajima’s test statistics as described by Nei M and Kumar [29] and by Tajima F. [30]. Tajima’s (D) test was 4.149 and nucleotide diversity(π) 0.603 P< 0.05 at 269 segregating sites for the 2 populations. For Malindi Tajima’s D test was 3.822 and nucleotide diversity of 0.654 P< 1.0 and for Tana River D= 1.446 and(π) was 0.318 p < 0.05.

### Tajima’s Relative Rate Test

This was done to test the equality of evolutionary rate between sequences A (ML4-2002) and B (ML5-ML11) with sequence C (TR4-2011) as the out-group. Chi-square (X^2^) test was 9.32 (P=0.00226, degree of freedom=1, P – value less than 0.05 is often used to reject the null hypothesis of equal rates between lineages. The analysis involved 3 nucleotides sequences Codon positions included were 1^st^ +2^nd^ +3^rd^ + Non-coding. On testing the relative evolutionary rate between all other sequences the P values were less than 0.05 and this made us conclude that, there were differences in lineages of all the sequences. All positions containing gaps and missing data were eliminated. There were a total of 285 positions in the final dataset. Evolutionary analyses were conducted using MEGA7 [27].

## DISCUSSION

### Molecular and genetic analysis of Malindi and Tana River Delta *Wuchereria bancrofti* Isolates in Kenya

Monitoring infection rates in human population and vectors is an essential component of any lymphatic filariasis control Programme: for identifying endemic areas where MDA is needed, for determining the progress of the program for deciding when to stop MDA and for the certification of elimination of the disease. Here, we provide information on genetic characterization of *Wuchereria bancrofti* in 2 endemic regions in Coastal Kenya. In this study, 18S rRNA gene was amplified, sequenced and analyzed. The amplified samples were collected from Malindi in 2002 just before the start of MDA in 2003 and in Tana River in 2008 before the first MDA; this was essential for the baseline data to enable us track any genetic changes in the isolates. The blasted sequences had an identity of 79-98% and this showed that there was relationship of the Kenya strains and other related parasites from other areas in NCBI data base.

Understanding the genetic differences in *Wuchereria bancrofti* could provide insight into effectiveness of drug regimes, the optimal time-course of drug administration and the potential of development of drug resistance. The nucleotide amplification, sequencing and phylogenetic reconstruction of the Ssp 1 DNA repeat sequence in this study revealed that there are genetic variations in *W. bancrofti* found in different geographical areas in Kenyan endemic regions. Despite the fact that this study relied on the available samples which were not of consecutive years of MDA, the overall genetic divergence observed for the entire population was 1.11. This information on genetic variability of *W. bancrofti* in Kenyan *W. bancrofti* population added to the very few studies that aimed at understanding the genetic heterogeneity of *W. bancrofti* [19], [31], [32]. These variations could be attributed to drug pressure and also geographical distribution in endemic regions of Kenya. These findings were in agreement with the results obtained De-Souza and colleagues [33] who found a considerable genetic variability within *W. bancrofti* populations in Ghana. There was a great divergence observed within Malindi population (0.95) compared to that of Tana River Delta (0.4), this can be attributed to the long period Malindi has been on MDA (from 2003) to Tana River (from 2011). By 2015 Malindi had received a total of 5 rounds while Tana River had received only 1 rounds of MDA [12]. Although a combined single dose of Albendazole and Diethylcarbamizine or Ivermectin has been used by Lymphatic filariasis control programmes effectively, reliance on these drugs only is highly vulnerable to emergence and spread of drug resistance. This might be anticipated due to drug pressure leading to intensive and prolonged new selection pressure on the parasites, which may in turn affect the genotypic and phenotypic structure of a parasite population [33], [34], [35]. The genetic differences observed in this study may also be attributed to environmental selection pressures and may explain the epidemiological findings of *W. bancrofti* distribution in endemic in Kenya. Taking this in to account, the emergency of new W. b resistant strains against the current choice of drug cannot be ruled out. These variations observed with the Kenyan strain could lead to drug resistance by different parasites strains.

A significant decline of *W. bancrofti* prevalence in Kenya has been noted [10], [11], [12] following MDA implementation and vector control programmes in Kenya. It is however evidenced MDA may lead that to genetic variability or resistance strains of *W. bancrofti* and this may result to resurgence of disease after elimination or increased prevalence. For instance, it is reported in some other endemic countries such as the Polynesian Islands of Moorea and Maupiti, over 50 years of MDA using DEC did not eliminate the disease [36]. It is presumed that the genetic differences in parasite population could be one of many possible explanations for this failure. These differences could be an indicator of selection pressure on the parasite suggesting that perhaps environmental selection pressures or other factors are in play. Though known cases of LF associated drug resistance in humans have not yet been reported, the development of drug resistance is a realistic possibility as drug resistance to both Ivermectin and Albendazole is prevalent in nematodes of veterinary importance [37]. Thus, an understanding of the genetic distinctness of various parasite populations could be a useful indicator in assessing and responding to the development of drug resistance in the future. A study by Schwab and colleagues [37] utilized populations of *W. bancrofti* infected individuals treated with 400mg albendazole in combination with 600 mg/kg Diethylcarbamizine and untreated areas in Burkina Faso and Ghana and observed resistance allele at 26.2% frequency in untreated populations, 60.2% in populations treated once with one drug and 86.2% in populations treated twice with the drug combination. He also observed that in the 2 untreated Ghanaian populations, the resistant allele was at a frequency of 2.7% and 0.33%. Also 2 different genetic variants of the parasite have been reported, with high genetic divergence and gene flow in different geo-climatic regions in India [38], [39]. Generally, genetic variability in *W. bancrofti* parasites may affect the success of MDA programs despite the fact that current elimination program assumes no differences within the parasite population and same treatment is administered to at risk population [40]. The finding on the genetic heterogeneity of the populations at different places within microfilaria carriers therefore calls for appropriate chemotherapeutic strategies for the elimination of lymphatic filariasis [31], [41].

To determine genetic divergence between species and within sub-species, genetic distance was evaluated. Species with closer genetic relationship have smaller genetic distance as compared to species with larger genetic distance. In this study, within host genetic structure was evaluated by comparing genetic distances both within and between the 2 populations. It should however be noted that Malindi is a cosmopolitan area and human and vector migration may significantly have influenced population structure of *W. bancrofti*. The fact that *W. bancrofti* undergoes development in both human and vectors, transmission dynamics isolation by distance may be due to movement of infected people, vector dispersal or combination of these factors.

Phylogenetic tree construction was done to assess the relatedness among sequences of *W. bancrofti* isolates of this study and other related sequences retrieved from the gene bank. Preliminary studies by Rekha and colleagues [42] demonstrated that the *W. bancrofti* population exhibited a trend of clustering according to drug treatment. This has been exhibited in our current study: Tana River Delta isolates of 2008 and 2011 samples clustered together which also indicated that there was no much divergence in Tana River isolates. The 8 Malindi isolates clustered in to 2 clusters and this indicates some genetic variability. Isolates of the same year from Malindi (2002) did not cluster together indicating some differences in the isolation villages probably by distance. Closely related were Malindi isolates labeled ML1 and ML4 and these were isolates from the same village in Malindi during the same year 2002, indicating that there were no much different in strains of the same geographical area just before treatments begun (Figure 4). The Malindi isolates first cluster and the Tana River Delta Clusters have a relationship with *Brugia pahangi* ribosomal protein S13, *W. bancrofti* ribosomal protein S13 and *B. Pahangi* mRNA accession numbers M86643.1, M86642.1 and X16591.1 respectively as per the reconstructed rooted phylogenic tree (figure 4). The common origin of the Kenya Isolates both from Tana River Delta and Malindi was *W. bancrofti* isolates W. b 1-07 18s rRNA accession number EU272178.1, from Brazil. In the analysis, *Loa loa* 18s rRNA accession number DQ995497.1from Spain was used as the out-group.

The Neutrality and selection test results showed that strong selection was occurring in *Wuchereria bancrofti* populations in the 2 Kenyan endemic areas. The two populations had a positive Tajima’s D value which reveals that mutations have resulted in high nucleotide diversity (0.603) with a P-value of less than 0.05 (p < 0.05). For these results null hypothesis of equal rates between lineages is rejected, and thus in our populations, the isolates had different rates of lineages and thus mutation was observed. Malindi isolates had a higher mutation rate compared to Tana River Delta isolates. There is a possibility of resistant strains developing from Malindi strains as compared to Tana River Delta.

## Conclusion and Recommendations

The analysis of genetic profiles of *W. bancrofti* from two endemic regions of Kenya indicated an existence of considerable genetic variability among parasite populations. In our study, *W. bancrofti* isolates of Malindi region were highly variable compared to that of Tana River Delta isolates. This could have been due to evolution of the parasites owing to the drug stress, environmental variations, infection transmission from other areas by human migration and parasite evolution to overcome the MDA drugs. The data adds to our understanding of the phylogenic diversity of these devastating parasites and the genetic information could support the control and monitoring of LF in these endemic areas. Further study in genetic variation aspects will shed light on the specific factors responsible for such divergence.

Further studies to better understand the population structure and genetic differentiation of this parasite will provide important insights into patterns of transmission, disease outcome, and anthelmintic drug resistance, and influence the design and implementation of public health interventions aimed at eliminating this disease.

## Availability of data and materials

Data on the sequences can be found in the Genbank.

## Acknowledgement

In this paper I wish to acknowledge Jacinta Muli for her assistance in laboratory processes and data analysis.

## Consent for publication

This paper is published with the permission of the Director, Kenya Medical research Institute.

## Roles of Investigators

**KNM** Conceived and developed the proposal, sourced for the funding, carried out laboratory experiments data analysis and developed the Manuscript. **JK** reviewed the proposal, provided guidance throughout the study, provided the archived specimen, and manuscript development and review. **WL, MW, MC, KL** helped in the proposal development, study design, supervised the study and reviewed the manuscript. **LS, WD, IC, WE** designed the study, reviewed the proposal, helped in data analysis and manuscript development and review. **GR, LJ** Reviewed the proposal, study design, helped in laboratory analysis and review of the manuscript.

